# Pre-meiotic, 21-nucleotide Reproductive PhasiRNAs Emerged in Seed Plants and Diversified in Flowering Plants

**DOI:** 10.1101/2020.10.16.341925

**Authors:** Suresh Pokhrel, Kun Huang, Sébastien Bélanger, Jeffrey L. Caplan, Elena M. Kramer, Blake C. Meyers

**Affiliations:** Donald Danforth Plant Science Center, Saint Louis, MO 63132; Division of Plant Sciences, University of Missouri-Columbia, Columbia, MO 65211; Bio-Imaging Center, Delaware Biotechnology Institute, University of Delaware, Newark, DE 19711, USA; Department of Organismic and Evolutionary Biology, Harvard University, Cambridge, MA, 02138

## Abstract

Plant small RNAs (sRNAs) are important regulatory elements that fine-tune gene expression and maintain genome integrity by silencing transposons. They have critical roles in most pathways involved in plant growth and reproductive development. Reproductive organs of monocots produce abundant phased, small interfering RNAs (phasiRNAs). The 21-nt reproductive phasiRNAs triggered by miR2118 are highly enriched in pre-meiotic anthers, and have not been described in eudicots. The 24-nt reproductive phasiRNAs are triggered by miR2275, and are highly enriched during meiosis in many angiosperms. Here, we describe additional variants of 21-nt reproductive phasiRNAs, including those triggered by miR11308 in wild strawberry, a eudicot, and we validate the presence of this pathway in rose. We report the widespread presence of the 21-nt reproductive phasiRNA pathway in eudicots, with novel biogenesis triggers in the basal eudicot columbine and the rosid flax. In eudicots, these 21-nt phasiRNAs are enriched in pre-meiotic stages, a spatiotemporal distribution consistent with that of monocots and suggesting a role in anther development. Although this pathway is apparently absent in well-studied eudicot families including the Brassicaceae, Solanaceae and Fabaceae, our work in eudicots supports a singular finding in spruce, indicating that the pathway of 21-nt reproductive phasiRNAs emerged in seed plants and was lost in some lineages.

## Introduction

PhasiRNAs are generally produced by the action of 22 nucleotide (nt) miRNA triggers on polyadenylated messenger RNA (mRNA) or long, non-coding RNA (lncRNA) targets generated by RNA Polymerase II^1^. These target transcripts are subsequently processed by the action of RNA-DEPENDENT RNA POLYMERASE 6 (RDR6) and DICER-LIKE (DCL) proteins into duplexes of 21- or 24-nt siRNAs. These siRNAs are “in phase” with one another; that is, the siRNAs map to the genome with regular spacing, the result of processive cleavage by Dicer from a long precursor (a PHAS precursor). The best characterized phasiRNAs in plants are trans-acting siRNAs (tasiRNAs) which function in development by regulation of auxin signaling ^1^.

Reproductive tissues of grasses contain both 21- and 24-nt phasiRNAs, derived from non-coding precursors encoded at hundreds of genomic loci. The 21-nt phasiRNAs are triggered by miR2118, and are mainly enriched in pre-meiotic anther tissues, during the stages at which the specification of cell fate occurs; they originate in the epidermal layer of the anther, but accumulate subepidermally ^2^. The 24-nt phasiRNAs are abundant in the meiotic stages of anther tissues and are largely triggered by miR2275, although in some monocots, the miR2275 trigger is absent^3^. The spatiotemporal pattern of 21-nt phasiRNAs in male reproductive tissues has been well-described in maize and rice^2,4^. These reproductive, 21-nt phasiRNAs lack obvious or at least validated targets, but play a role in photoperiod sensitive male sterility in rice^5^. A mutant in rice of an Argonaute protein that selectively binds these 21-nt phasiRNAs^6^ is male sterile. These data demonstrate that the 21-nt reproductive pre-meiotic phasiRNAs and/or their functions are required for male fertility, and thus they are hypothesized to function in some important aspect of reproductive development.

The evolutionary origins of 21-nt reproductive phasiRNAs are poorly examined. Their presence in monocots is clear, as the 21-nt, pre-meiotic phasiRNAs were first reported in grasses^7^ and later described from just three loci in asparagus but still enriched in the middle layer, tapetum, and archesporial cells^3^. Despite extensive analyses, 21-nt reproductive phasiRNAs have not been reported in several well-studied plant families, including the Brassicaceae, Fabaceae and Solanaceae, leading to our own prior assumption that they are absent from the eudicots. In earlier work from our lab, we described a set of 21 noncoding loci in Norway spruce that are targets of miR2118, produce 21-nt phasiRNAs and are enriched in abundance in male cones^8^. This observation, as-yet unsupported or reproduced from analyses of non-monocot angiosperms, indicated the emergence of the 21-nt reproductive phasiRNAs outside of the monocots, in gymnosperms. Here, we describe the discovery of the pathway of pre-meiotic, 21-nt, reproductive phasiRNAs in several eudicots: wild strawberry (*Fragaria vesca*), rose (*Rosa chinensis*), the basal eudicot columbine (*Aquilegia coerulea*), and flax (*Linum usitatissimum*). We conclude that, like the 24-nt reproductive phasiRNAs^9^, the 21-nt reproductive pathway is widespread in angiosperms and may even, with origins in seed plants, may have emerged prior to the 24-nt reproductive phasiRNAs.

## Results

### Pre-meiotic anthers of wild strawberry produce abundant 21-nt phasiRNAs

We analyzed sRNAs in tissues of wild (diploid) strawberry (*F. vesca*), as this plant was highly informative for our earlier analyses of 24-nt, reproductive phasiRNAs^9^; in fact, we were considering it as possible model system for the study of 24-nt phasiRNAs. We characterized 21-nt phasiRNAs in both vegetative and reproductive tissues of wild strawberry, and we identified 25 loci that give rise to 21-nt phasiRNAs (“21-*PHAS*” loci) that were abundant specifically at anther stage 7 (Fig. 1a). This stage corresponds to the pre-meiotic stage of anther development^10^. These unannotated loci were mostly predicted to be non-coding (Supplementary Data 1); they contained only one conserved sequence motif, which is the target site of miR11308-5p (henceforth, miR11308) (Fig. 1b, 1c). miR11308 has three mature variants (Fig. 1b, 1d), two of which derive in the genome from polycistronic precursors (Supplementary Fig. 1A, Supplementary Data 2). Mature miR11308 accumulation peaked at anther stage 7 (Fig. 1b), similar to the peak of the reproductive-enriched, 21-nt phasiRNAs. For these loci, the most abundant, phased 21-nt phasiRNA is generated from the cleavage site of miR11308 (or in a register spaced by 21 bp), as exemplified in Fig. 1e. These loci were distributed across all but chromosomes 2 and 3 of the *F. vesca* genome, while miR11308 originated from chromosome 6 (Supplementary Fig. 1B). We were surprised to find pre-meiotic, 21-nt reproductive phasiRNAs, as we could find no record of a previous report of their presence in a eudicot.

**Figure 1.**
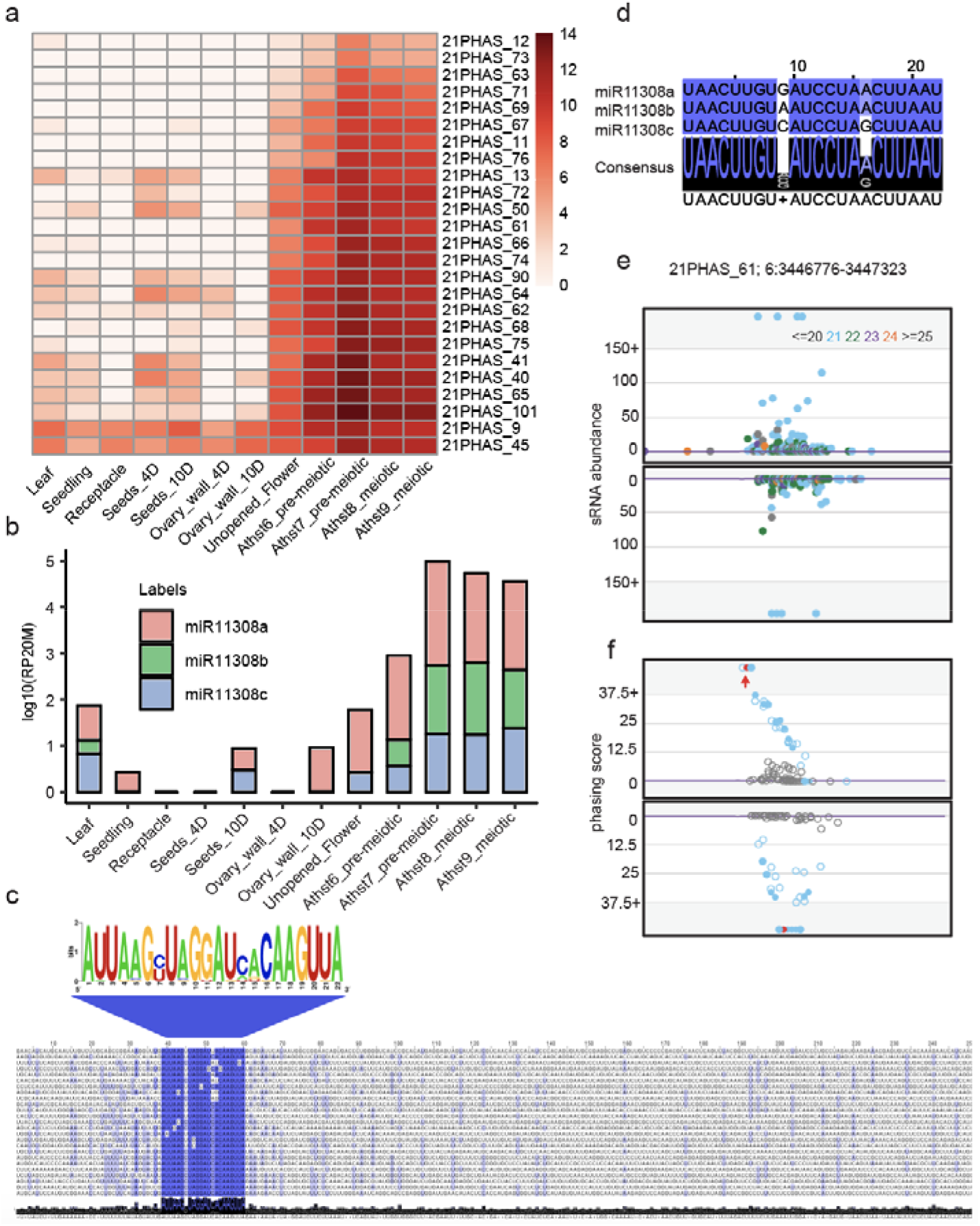
Reproductive 21-nt phasiRNAs triggered by miRNA miR11308 are abundant in wild strawberry a. Expression of 21-nt reproductive phasiRNAs in different tissues and anther development stages. The key at right indicates the abundance in units of log2(RP20M). Athst: anther stage. RP20M: reads per twenty millions mapped reads. b. Abundance of miR11308 members in log10(RP20M) in different tissues of wild strawberry. c. Above: Sequence logo denoting conservation of target site of miR11308 for 25 21-*PHAS* loci. Below: Nucleotide sequence alignment of 21-*PHAS* loci with sequence similarity denoted by the intensity of the blue color showing that the miR11308 target site is the only conserved region for all the precursors. d. Alignment of members of the miR11308 family in wild strawberry. The degree of conservation is denoted by intensity of the blue color; the consensus sequence of the alignment is shown with a sequence logo. e. Abundance (RP15M) of small RNAs in both strands of an example locus padded with 500 base pairs, each side. sRNA sizes are denoted by colors, as indicated at top. f. Phasing score of same locus as (e); the red dot indicates the highest phased sRNA position. The red dot also represents the coordinate (3446776) which exactly coincides with the predicted cleavage site of miR11308, denoted by the red arrow.

In maize and rice, mature variants of the miR2118 family trigger production of the 21-nt reproductive phasiRNAs^2,4^. In many other species, the miR482 family, a predecessor and relative of the miR2118 family, triggers phasiRNAs from disease resistance genes^11^. However, in wild strawberry, only six miR2118/miR482-derived non-coding 21-*PHAS* loci are enriched during anther stages (Supplementary Data 1). We found 76 21-*PHAS* loci triggered by miR2118/miR482 in total; 65 loci are abundant in vegetative tissues while 11 loci are enriched in reproductive tissues. These loci are mostly from protein coding genes, and are mainly similar to genes encoding disease resistance proteins, mirroring the 21-*PHAS* loci of other eudicots, such as soybean and Medicago^12,13^ (Supplementary Fig. 2A, Supplementary Data 1). In strawberry, we found four precursors give rise to 15 mature variants of miR2118/miR482; these variants accumulate in all tissue types, with greater abundances found in seeds (Supplementary Fig. 2B and C). Therefore, unlike grasses, in wild strawberry, the majority of 21-nt reproductive phasiRNAs are triggered not by miR2118 but rather by a lineage-specific miRNA, miR11308.

We next asked where this trigger of 21-nt reproductive phasiRNAs localizes. *In situ* hybridization localization of miR11308 in wild strawberry showed that it localizes to microspore mother cells (MMCs), meiocytes and tapetal cells (Fig. 2a, 2b) unlike the miR2118 in maize which localizes only in epidermis^2^. miR11308 is more abundant in MMCs of pre-meiotic cells and in tapetal cells of meiotic stages compared to the post-meiotic stage. Next, we examined the localization patterns of the most abundant 21-nt phasiRNAs from these loci with smFISH (Supplementary Fig. 3A) using a pool of fluorescently-labeled probes against 50 21-nt phasiRNAs. Surprisingly, we found these molecules were localized in all cell layers, most abundant in the MMC during the pre-meiotic stage of anther development (Fig. 3a). By the tetrad stage, the phasiRNAs are only detectable in the MMCs, and barely detectable in other cell layers. This spatiotemporal distribution of 21-nt phasiRNAs is generally similar to the accumulation pattern found in grasses^2,14^and thus these 21-nt phasiRNAs in strawberry may play a role in male fertility.

**Figure 2.**
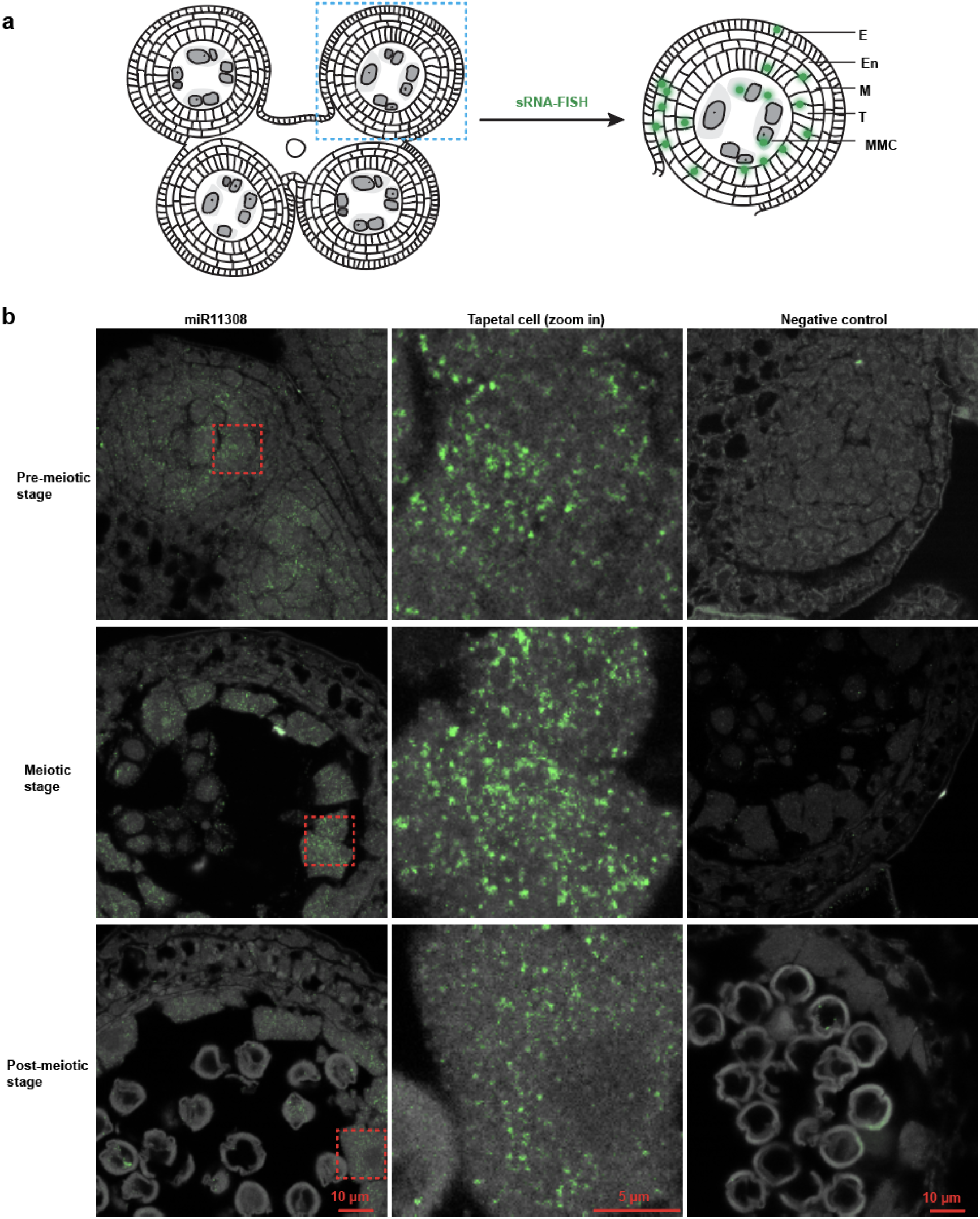
miR11308 accumulates in microspore mother cells, meiocytes, and tapetal cells in the anthers of wild strawberry a. Schematic diagram of the sRNA-FISH method in anther stages shown in panel b. E: epidermis, En: endodermis, M: middle layer, T: tapetum, MMC: microspore mother cell. b. In situ hybridization of miR11308 in pre-meiotic, meiotic and post-meiotic anther stages

### Conservation of miR11308 and 21-nt reproductive phasiRNAs in other Rosaceae species

To determine whether this pathway is conserved in other members of the Rosaceae family and beyond, we investigated the presence of the miR11308 in species of Rosaceae and Betulaceae for which a genome sequence is available. We found that the pattern of miR11308 presence is consistent with its emergence in the Rosaceae subfamily Rosoideae, as it is present in at least four genera (Fig. 3b, Supplementary Data 2). Similar to miR2275^9^, this miRNA generates mature sequences with a polycistronic precursor cluster which itself is conserved in the Rosoideae. We hypothesize that this polycistronic nature of the precursors is of functional importance for the biogenesis of reproductive phasiRNAs.

**Figure 3.**
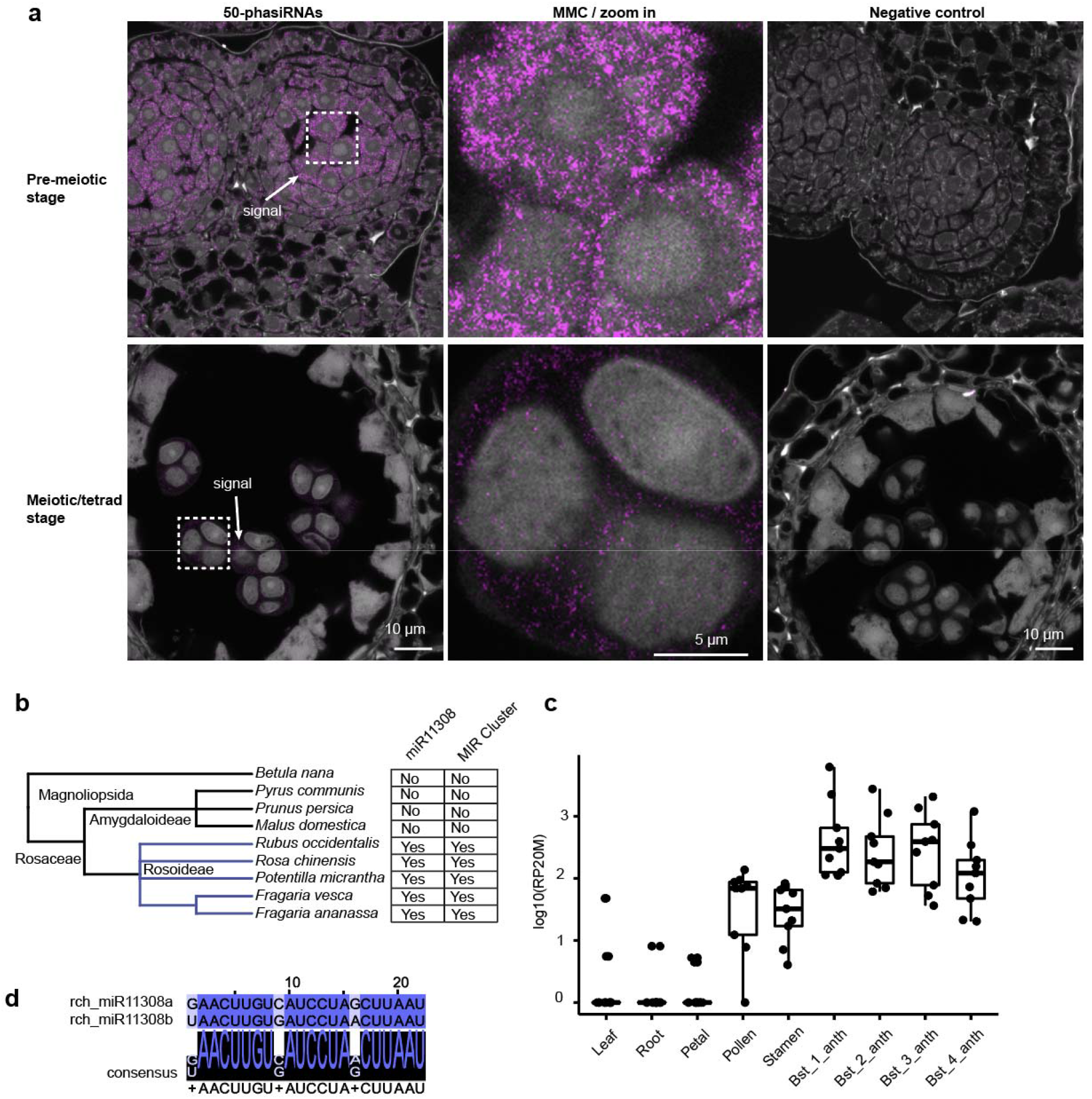
21-nt reproductive phasiRNAs localize in all cell layers of the pre-meiotic strawberry anthers and the miR11308 trigger emerged in the Rosoideae. a. Localization of abundant 50 21-nt reproductive phasiRNAs in pre-meiotic and meiotic anther tissues of wild strawberry. b. Phylogenetic tree showing conservation of miR11308 in the Rosaceae. c. Abundances of summed 21-nt phasiRNAs in different tissues of rose in units of log10(RP20M). In the boxplot, the center line represents the median, box limits are the upper and lower quartiles; whiskers are the 1.5x interquartile range; points show the scatter of data points for nine 21-*PHAS* loci. Bst indicates bud stage, anth indicates anther. d. Alignment of members of miR11308 in rose.

We next analyzed the small RNA transcriptomes of rose and identified nine 21-*PHAS* loci enriched in the anthers of early reproductive bud stages (Fig. 3c and Supplementary Fig. 3B,), thus confirming the presence of 21-nt reproductive phasiRNAs in this species. Similar to wild strawberry, these loci are mostly non-coding (Supplementary Data 3) and are also targeted by two variants of miR11308 (Fig. 3d). We further found that three precursors of six mature variants of the rose miR2118/482 family target 86 21-*PHAS* loci; we found them to be abundant in all tissue types, and mostly derive from genes coding for nucleotide binding–leucine-rich repeat proteins otherwise known as “NLRs” that commonly function as innate immune receptors (Supplementary Fig. 3C and D, Supplementary Data 3). Hence, we report that the miR31108 has acquired a newly identified role to generate 21-nt reproductive phasiRNAs in the Rosaceae subfamily Rosoideae.

### Novel trigger miRNAs and conservation of 21-nt reproductive phasiRNAs in the basal eudicot columbine and in flax

We next examined the small RNA transcriptomes of vegetative tissues and four different bud stages (Supplementary Fig. 4A) of the basal eudicot *Aquilegia*, (common name ‘columbine’) to determine whether 21-nt reproductive phasiRNAs are present and/or conserved in this species. We identified 112 21-*PHAS* loci; 91 enriched in reproductive tissues (Fig. 4a) and an additional 21 21-*PHAS* loci that were expressed similarly in all tissue types (Supplementary Fig. 4B). Out of 112 loci, we found that 65 loci are targeted by a new miRNA, aco-cand81; most loci (63/65) are from non-coding regions of the genome (Supplementary Data 4), and they possess a conserved 22 nt motif: the target site of aco-cand81 (Supplementary Fig. 4C). The aco-cand81 miRNA has six mature variants from seven different precursors (Fig. 4b, Supplementary Data 2), with one polycistronic cluster generating two mature variants (Supplementary Fig. 4D). The accumulation pattern of aco-cand81 is similar to the 21-nt phasiRNAs initiated by this miRNA (Fig. 4a, b).

**Figure 4.**
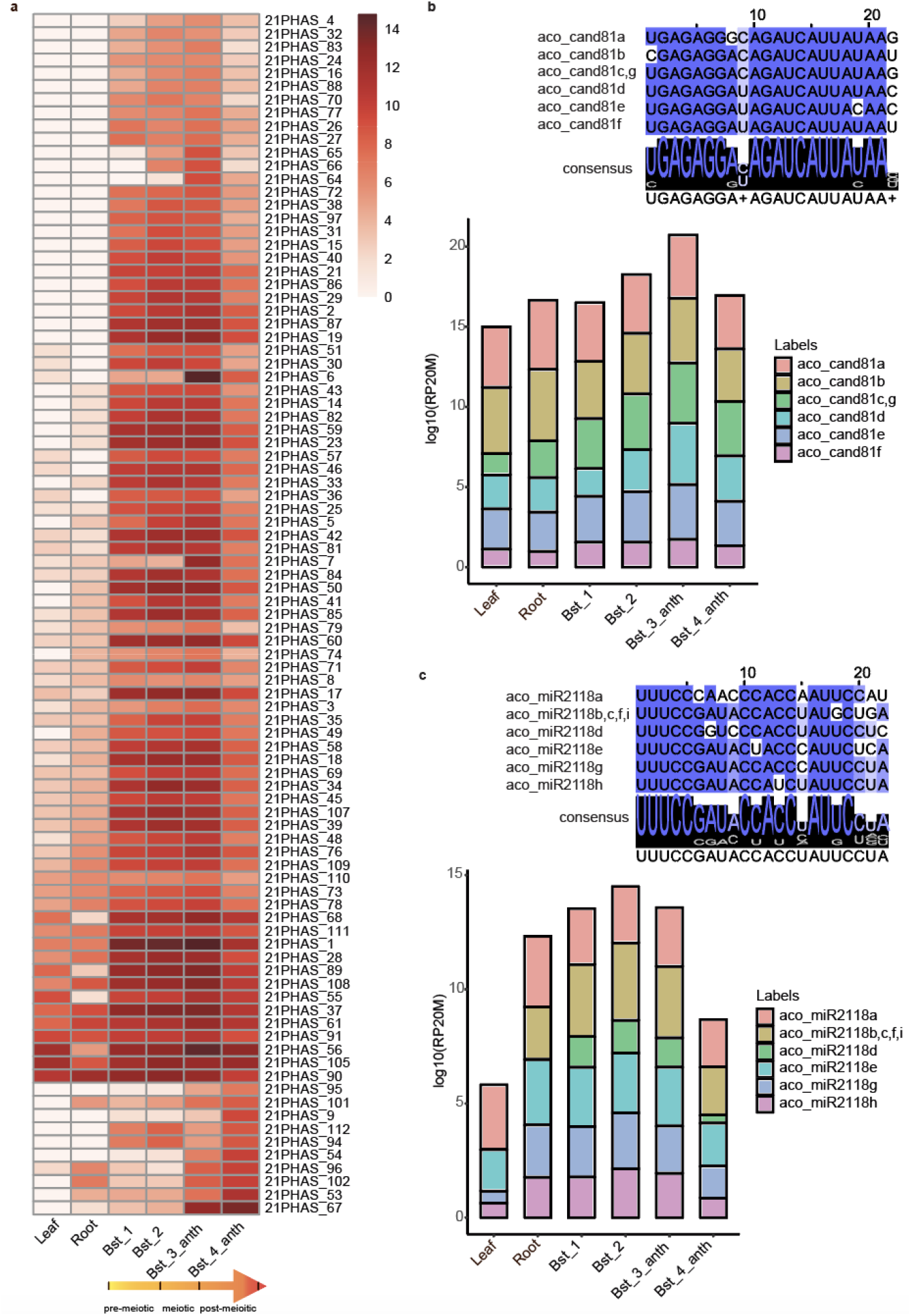
Conservation of reproductive 21-nt phasiRNAs and their triggers in the basal eudicot columbine. a. Expression of 21-nt reproductive phasiRNAs in different vegetative tissues and anther development stages in columbine. The lowermost ten loci are most enriched in stage 4.The yellow gradient arrow indicates the developmental stages of anthers. The key at right indicates the abundance in units of log2(RP20M). Bst indicates bud stage, anth indicates anther. b. Above: Alignment of members of aco_cand81 in columbine. The degree of conservation is denoted by the intensity of the blue color and the consensus sequence of the alignment is shown in a sequence logo. Below: Abundance of aco_cand81 members in log10(RP20M) in different tissues of columbine. c. Similar to panel b but for the miR2118/482 family in columbine.

The remaining 47 (of 112) 21-*PHAS* loci are targeted by mature variants of the miR2118/482 family (Fig. 4a, c and Supplementary Fig. 5A). Among these, 37 loci are abundant in bud stage 2 and 3 (Fig. 4a) and are non-coding, whereas the other 10 loci are abundant in all tissues type and are mostly derived from NLR protein-coding genes (Supplementary Fig. 4A and Supplementary Data 4). The miR2118/482 family has six mature variants that are derived from nine different precursors with two polycistronic clusters (Fig. 4c, Supplementary Fig. 5B, and Supplementary Data 2). The miR2118/482 family members are highly abundant at bud stages 2 and 3, similar to the phasiRNAs they trigger (Fig. 4c). Most of the reproductive 21-*PHAS* loci in *Aquilegia* are concentrated on chromosome 4, an unusual chromosome showing higher polymorphism compared to other chromosomes between the ten different *Aquilegia* species^15^. The aco-cand81 miRNA precursors were derived from chromosomes 4 and 1 whereas miR2118/482 precursors were distributed on chromosomes 1, 3, and 5 (Supplementary Fig. 6). Unlike grasses, in *Aquilegia*, 21-nt reproductive phasiRNAs are triggered by two miRNA family members, where at least one precursor is polycistronic.

To more precisely identify the developmental stages of the buds/anthers that we sequenced, anthers from seven bud lengths were examined. We found anthers from 2 to 4 mm buds are in pre-meiotic stages while 5 to 6 and 10 to 20 mm buds are meiotic and post-meiotic stages (Supplementary Fig. 7). Therefore, the 21-nt phasiRNAs in columbine initiate in pre-meiotic stages and their accumulation is maintained up to post-meiotic stages.

To further confirm the conservation of the 21-nt reproductive phasiRNA pathway in eudicots, we profiled small RNAs from both vegetative and reproductive tissues of flax, a species in the phylogenetic tree of eudicots outside the Rosaceae but within the rosids. We found 11 reproductive-enriched 21-*PHAS* non-coding loci triggered by miR2118/482 variants (Supplementary Fig. 8A, B and C) in flax (Supplementary Data 2 and 5). Unlike columbine, in flax, these 21-*PHAS* loci are triggered only by the canonical trigger of pre-meiotic phasiRNAs, as described from extensive work in monocots^2,4^ (miR2118/miR482).

### Genes important to the 21-nt reproductive phasiRNA pathway

Factors known to function in phasiRNA biogenesis are mainly RDR6, Argonautes (AGO), and DCL proteins. We analyzed presence of these proteins in total of 13 species: four eudicots (wild strawberry, rose, columbine, flax) we analyzed for phasiRNAs, two gymnosperms (Norway spruce, ginko), and other seven representative species for which these genes were already characterized: *Amborella*, soybean, tomato, *Arabidopsis*, *Asparagus*, maize, and rice (Fig. 5a, Supplementary Fig. 9A and B). RDR6 is responsible for making the double-stranded RNA precursors after miRNA cleavage during phasiRNA biogenesis and it is conserved in seed plants (Supplementary Fig. 9A). Among 13 species, RDR6 is present in two or more copies in all species except in rice, *Asparagus*, *Arabidopsis*, and columbine. Three eudicot species we analyzed for phasiRNAs have two *RDR6* copies but not columbine (Supplementary Fig. 9A). We hypothesize that there is a species-specific duplication of RDR6, potentially facilitating a sub-functionalization of their roles, as both copies are enriched in reproductive tissues (Supplementary Data 6). 21-nt phasiRNAs are produced by *DCL4*, which is conserved in seed plants (Supplementary Fig. 9B) and mostly enriched in reproductive tissues, except in rose (Supplementary Data 6). DCL3 and DCL5 are Dicer proteins responsible for the production of 24-nt siRNAs, with DCL5 specialized for 24-nt phasiRNAs in monocots and DCL3 hypothesized to functioning in a dual roles, producing 24-nt phasiRNAs in eudicots in addition to its well-described role in making heterochromatic siRNAs ^2,9^. There is a duplication of *DCL3* in all four eudicots genomes that we examined for phasiRNAs and in a gymnosperm ginko among all the species and duplicated *DCL3 (DCL3b)* was more enriched in reproductive tissues than *DCL3a* (Supplementary Data 6) suggesting the possibility of neo-functionalization of DCL3b for the production of 24-nt reproductive phasiRNAs in eudicots^9^.

**Figure 5.**
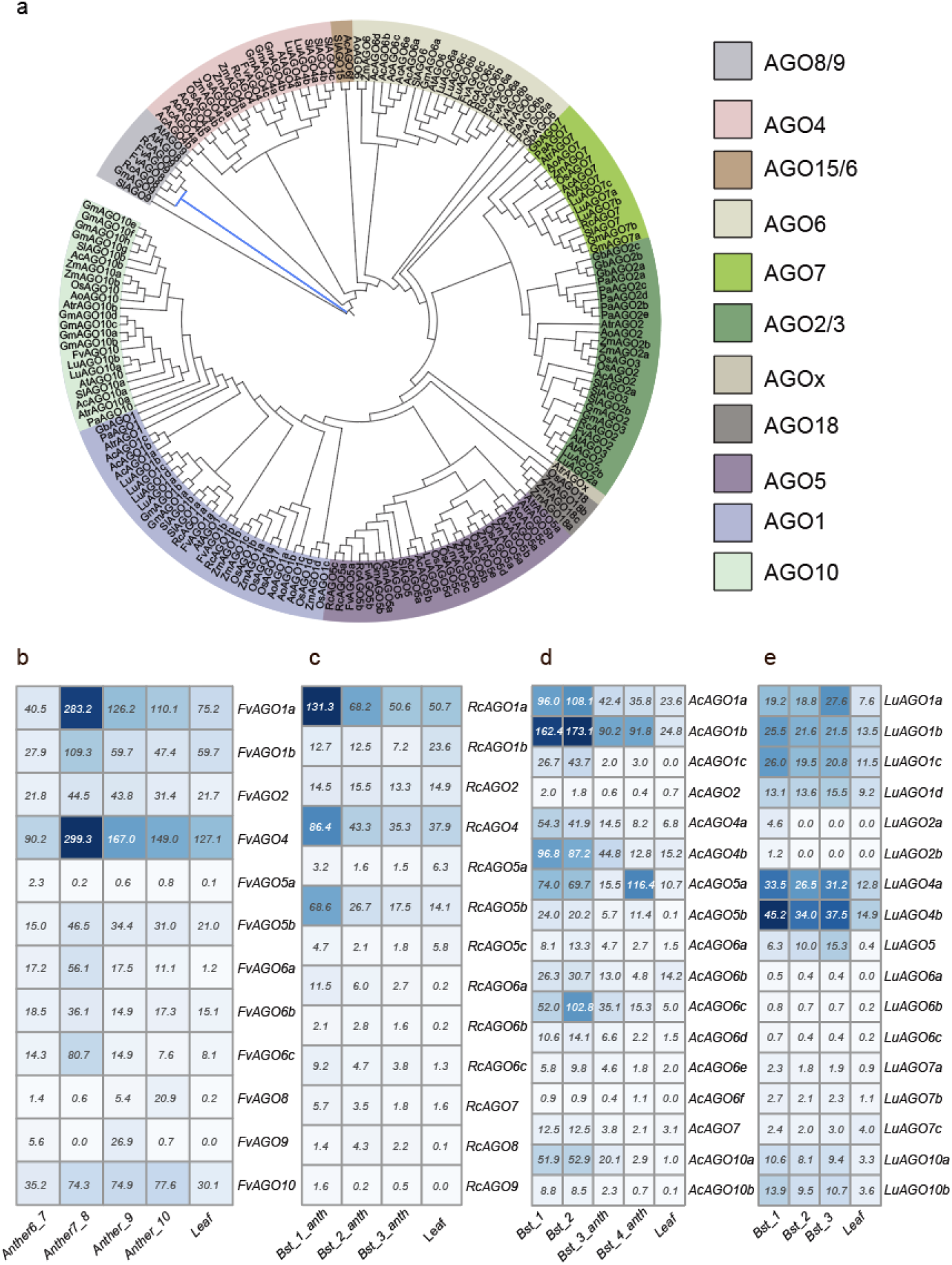
Characterization of AGO gene family members in wild strawberry, rose, columbine, flax and other representative species from gymnosperms, eudicots, and monocots. a. Phylogenetic tree of Argonaute protein family members encoded in the genomes of plant species, with two-letter prefix indicating the source genome: wild strawberry (Fv), rose (Rc), columbine (Ac), flax (Lu), Norway spruce (Pa), ginko (Gb) identified in this study along with other seven representative species: *Amborella* (Atr), soybean (Gm), tomato (Sl), *Arabidopsis* (At), *Asparagus* (Ao), maize (Zm), and rice (Os). Abundance profile of genes encoding AGO family members in (b) wild strawberry, (c) in rose, (d) in columbine, and (e) in flax. Bst indicates the bud stage, anth indicates the anther.

We identified 12, 13, and 17 AGO proteins encoded in the wild strawberry, rose, columbine and flax genomes, respectively (Fig. 5a). All of the identified AGO transcripts are expressed ≥ 0.5 TPM (transcripts per million) in these four species (Fig. 5b). In rice, AGO1b/1d were implicated in loading miR2118 and U-21-nt phasiRNAs^14^ while AGO5c (i.e., rice MEL1) is implicated for the loading of reproductive C-21-nt phasiRNAs^6,14^. Here, we identified two *AGO5* copies in strawberry, and both *AGO5* copies are enriched in pre-meiotic stages while in rose, among three *AGO5* copies, only *AGO5b* (orthologous to *AGO5b* in strawberry) was enriched in reproductive tissues. In flax, one copy of *AGO5* is enriched in reproductive tissues. In columbine both copies of *AGO5* are enriched in reproductive tissues but *AGO5b* was more enriched (Supplementary Data 6). Overall, based on phylogeny and their expression profiles, AGO5 or AGO5a/b are the candidate AGO proteins for loading pre-meiotic 21-nt phasiRNAs in eudicots, as they are in monocots. Among AGO1 family members, in columbine, *AGO1c* is more enriched in reproductive stages, while in flax, strawberry, and rose, *AGO1a* is more enriched in reproductive tissues (Fig. 5b, Supplementary Data 6). So, AGO1c for columbine, and AGO1a for other three eudicots might be the effector protein for triggers of 21-nt reproductive phasiRNAs. Orthologues of AGO18 are absent in eudicots, gymnosperms, *Amborella*, and *Asparagus* consistent with its hypothesized origin in grasses (Fig. 5a). Genes encoding members of the AGO6 clade (specifically *AGO6a*) were enriched more than *AGO4* in reproductive tissues while *AGO9* in rose and strawberry are expressed exclusively in reproductive tissues (Fig. 5b, Supplementary Data 6). It is known that AGO4, AGO6 or AGO9 are binding molecules for 24-nt heterochromatic siRNAs during RNA-directed DNA methylation (RdDM)^16^. Thus, based on the expression and enrichment, AGO9 and AGO6 may also act as effector molecule for reproductive 24-nt phasiRNAs in eudicots.

## Discussion

We found that 21-nt reproductive phasiRNAs are present in several eudicots: wild strawberry, rose, the basal eudicot columbine, and flax. Whereas previous studies have shown that 21-nt reproductive phasiRNAs exist in one gymnosperm^8^ and more widely in monocots^2,4,7,17,18^ (Fig. 6a), this pathway is has not been described in several well studied eudicot families including the Brassicaceae, Fabaceae, and Solanaceae^2,12,13^. Another class of 21- or 22-nt secondary siRNAs active in plant reproduction are the epigenetically activated siRNAs (easiRNAs) that accumulate in *Arabidopsis* during stages of pollen maturation, derived from activated transposable elements^19,20^. easiRNAs appear to play a role in male gamete production and possibly post-fertilization genome stability and seed viability^21^. Even though both phasiRNAs and easiRNAs are secondary siRNAs important for plant reproduction, there is no evidence suggesting that they are related classes of RNAs, nor is it evident that easiRNAs are found outside of *Arabidopsis*. We hypothesize that 21-nt reproductive phasiRNAs are a more fundamental and broadly conserved pathway which emerged in gymnosperms but were lost in some lineages due to accommodations or adaptations of development.

**Figure 6.**
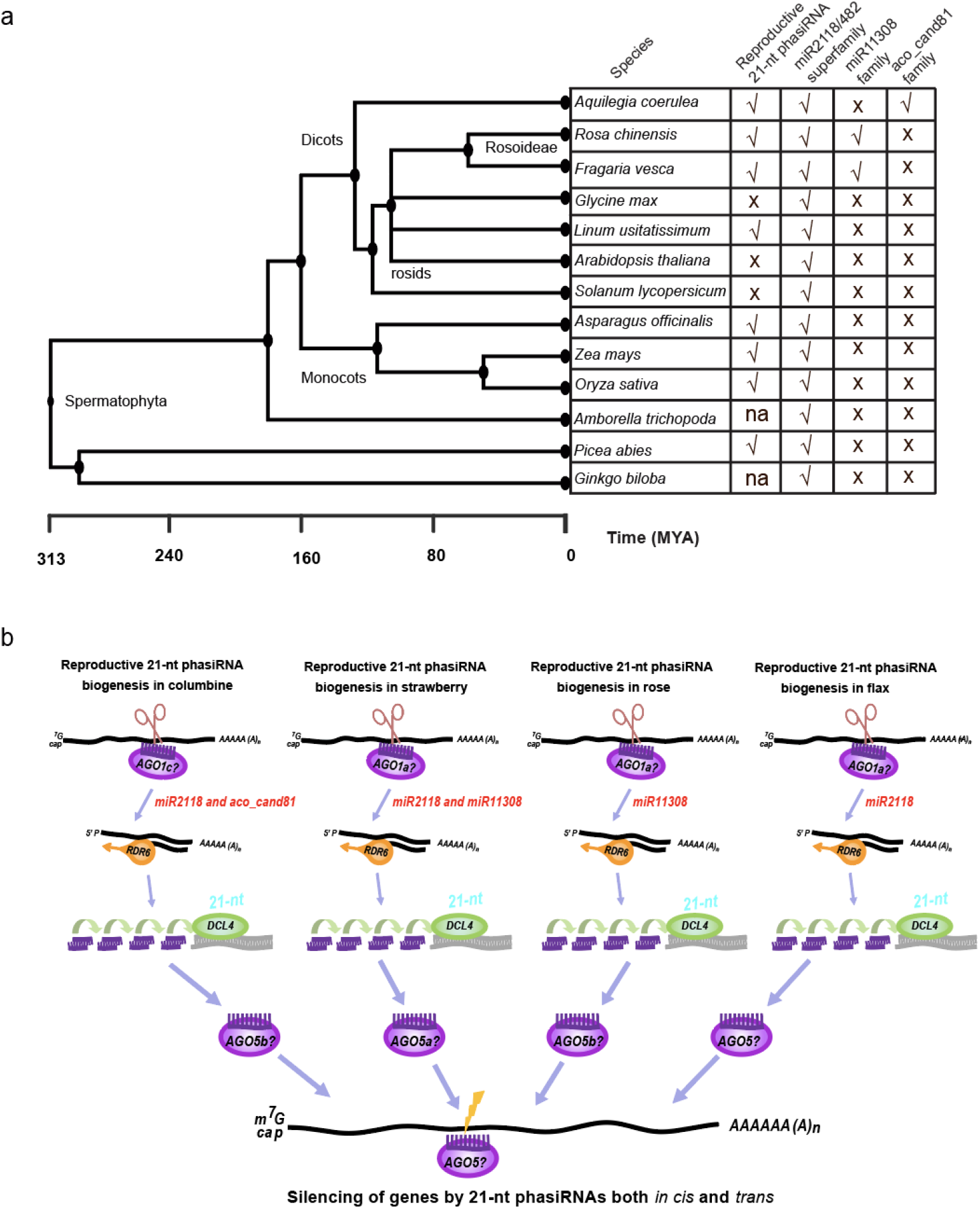
Conservation of reproductive 21-nt phasiRNA in seed plants. a. Phylogeny showing the representative species of seed plants according to estimated divergence times based on the time tree of life (Kumar et al., 2017). The miR2118/482 superfamily consists of miR2118, miR482, miR8558 and miR472. MYA means million years ago. √ indicates presence and X indicates absence; na indicates ‘not analyzed’. b. Model of generation of 21-nt reproductive phasiRNAs in seed and flowering plants. In gymnosperms, monocots, and some eudicots such as flax, miR2118 is the only trigger for the production of 21-nt reproductive phasiRNAs, while in columbine, rose and wild strawberry, there is a lineage-specific miRNA trigger for this role. Based on the roles in other species, family members of AGO1 and *AGO5* might load the trigger miRNA and 21-nt phasiRNAs respectively. The resulting phasiRNAs from these mechanisms were predicted to act in both *cis* and *trans* to modulate the gene expression.

In the basal eudicot columbine, we showed that reproductive tissues produce abundant phasiRNAs generated by both the canonical trigger miR2118/482 and a lineage-specific miRNA, aco-cand81, indicating a division of this phasiRNA biogenesis role. Moreover, the miRNA2118/482 family triggers 21-nt phasiRNAs from NLR genes in columbine, a conserved role of this miRNA from gymnosperms to angiosperms, particularly in eudicots^1^. The miR2118/482 derived *NLR* phasiRNA pathway also found in wild strawberry and rose, while the role of this miRNA family in generating reproductive 21-nt phasiRNAs is almost completely shifted to miR11308, a novel trigger of reproductive phasiRNAs; we also found evidence for the canonical trigger of reproductive phasiRNAs in wild strawberry, miR2118. In flax, we found that only miR2118/482 triggers the production of 21-nt reproductive phasiRNAs, consistent with a conserved role of the canonical trigger in this species. Based on all of these data, we propose a model of how different triggers in different species function to generate reproductive 21-nt phasiRNAs in seed and flowering plants (Fig. 6b). All of these triggers starts with uracil (U) and are 22-nt in length consistent with earlier finding that only 22-nt form of miRNAs in AGO1 complex are capable of initiating phasiRNA biogenesis^22^. Clearly more extensive data, from more diverse genomes of seed plants, would help improve the granularity of this model.

Similar to the 21-nt reproductive phasiRNAs in grasses, in wild strawberry, these phasiRNAs are also localized in all cell layers and abundant in MMCs of pre-meiotic anther suggesting similar role in reproductive development of eudicot anthers – whatever that role may be. Even more than a decade after describing the 21-nt reproductive phasiRNAs in rice and maize^7^, their precise molecular activities are unclear. Unlike miR2118 in grasses which localizes to the epidermis, miR11308 localizes in MMCs in pre-meiotic anthers, suggesting MMCs as origin of these pre-meiotic phasiRNAs. The localization of miR11308 in tapetal cells of meiotic anthers suggests other potential roles of this miRNA in later stages of anther development, yet to be examined.

In conclusion, the emergence and conservation of 21-nt reproductive phasiRNAs in eudicots supports an origin in the seed plants and suggests a functional role in reproduction. In rice, mutations in these 21-*PHAS* loci cause photoperiod-sensitive male sterility^5^, a key feature that has been leveraged for hybrid seed production. Therefore, the study of these 21-*PHAS* loci in a broader set of plant species may open up avenues to improve plant yields. Currently, it is unknown how these phasiRNAs modulate male fertility; future studies are needed to produce a deeper understanding of reproductive phasiRNAs. Some species may be better models than others, due to their life history, miRNA triggers, *PHAS* locus complexity, or other traits, as demonstrated by the diversity we observed in eudicots.

## Methods

### Plant material harvesting and RNA isolation

Vegetative tissues and unopened flower buds/anthers of columbine (cultivar, ‘origami’) were collected from greenhouse-grown plants under conditions of 20/18°C day/night and 16/8 hours light/dark. A total of four different bud stages (Supplementary Fig. 4A) were characterized, and buds were harvested for stage 1 and 2 and from stage 3 and 4, anthers were harvested. Two biological replicates were sequenced for each stage. Flax was grown in a growth chamber with 12h light at 25 °C followed by 12⍰h dark at 20⍰°C. Three bud stages (Supplementary Fig. 7A) were harvested and frozen immediately in liquid nitrogen. Rose anthers from four different bud stages (Supplementary Fig. 3B) were harvested and immediately frozen in liquid Nitrogen. Total RNA was isolated using the PureLink Plant RNA Reagent (ThermoFisher Scientific, cat. #12322012) following the manufacturer’s instructions. Total RNA quality and quantity were assessed by running gel and using qubit. Flax small RNAs (20 to 30 nt) were size-selected in a 15% polyacrylamide/urea gel and used for small RNA library preparation, while for other species direct total RNA was used for small RNA library construction. An aliquot of 5 μg of total RNA was used for size selection.

### RNA libraries and sequencing

Flax small RNA libraries were constructed using the NEBNext Small RNA Library Preparation set for Illumina (NEB, cat # E7300); for columbine and rose, RealSeq-AC miRNA Library kit for Illumina (Somagenics, cat # 500-00012) was used as per manufacturer’s recommendation. RNA-seq libraries were constructed using NEBNext Ultra II Directional RNA Prep kit for Illumina; RNA was treated with DNase I (NEB, cat # M0303S) and then cleaned using the RNA Clean & Concentrator™-5 (Zymo Research, cat # R1015S). Small RNA and RNA-seq libraries were single-end sequenced with 76 cycles. All libraries were sequenced on an Illumina Nextseq 500 instrument at the University of Delaware Sequencing and Genotyping Center at the Delaware Biotechnology Institute. For strawberry vegetative/reproductive tissues^9,23,24^ and rose’s vegetative tissues^25^, sRNA- and RNA-seq data were downloaded from public databases and used for the analysis.

### sRNA data analysis and PHAS annotation

Raw reads of small RNA libraries were processed by an in-house preprocessing pipeline^26^. PHASIS^27^ with a p-value cut off of 0.001 and ShortStack^28^ with default parameters were used to identify the PHAS loci in all of the species. PHAS loci from two different software were merged using BEDTools^29^. Only PHAS loci having target sites for miRNA triggers were annotated as valid PHAS loci. Target prediction was carried out by sPARTA^30^ with a target cut off score of ≤4. The small RNA abundance and phasing score were viewed at customized browser^31^. Based on the target site of triggers, 50 bp upstream and 500 bp downstream strand specific sequences were extracted from each PHAS loci and were annotated de novo by BLASTX^32^ against UniRef90. The coding potential of PHAS loci were assessed for protein coding potential by CPC (Coding Potential Calculator).

### RNA-seq data analysis and visualization

Raw reads were processed by an in-house preprocessing pipeline and Hisat2^34^ was used to align the reads with respective genomes using default parameters. Raw counts were obtained by using TPMCalculator^35^ and normalized to transcripts per million (TPM). To visualize the conservation of the target site of miRNA triggers, a multiple sequence alignment of PHAS loci strand-specific sequences was performed using MUSCLE^36^ with default parameters and we visualized the alignment using Jalview^37^. Weblogo images were created using logo generation form (http://weblogo.berkeley.edu/logo.cgi). Circular plots were made using OmicCircos^38^ for the chromosomal distributions, pheatmap (https://rdrr.io/cran/pheatmap/) and ggpubr (https://github.com/kassambara/ggpubr) were used to draw heatmaps and box-plot in R.

### Phylogenetic Analysis

Identification of RDR, Dicer and AGO families for all of the species were carried out by using Orthofinder^39^except for Norway spruce (*Pica abies*) for which genBlastG^40^ was used. The resulting protein sequences were visualized in CDvist^41^ to find complete domains. Curated proteins were aligned using default settings in PASTA^42^. The Maximum Likelihood (ML) gene tree for each protein family were generated using RAxML^43^ over 100 rapid bootstraps with options “-x 12345 ‒f a ‒p 13423 ‒m PROTGAMMAAUTO”. Trees were visualized and manipulated in iTOL^44^. The annotations used for the phylogenetic analysis for all 13 species for AGO, DCL and RDR proteins are listed in Supplementary Tables 7, 8 and 9 respectively.

### Tissue Embedding and Microscopy

Fresh anthers were dissected from different bud stages and fixed in a FAA solution overnight and dehydrated through a standard acetone series (30%, 50%, 70%, 80%, 90%, 100% of cold acetone) prior to being resin infiltrated and embedded using the Quetol (Electron Microscopy Sciences, cat #14640) using either heat polymerization. Embedded tissues were sectioned at 0.5 μm using the Leica Ultracut UCT (Leica Microsystems Inc.) and stained using a 0.5% Toluidine Blue O dye (, Electron Microscopy Sciences, cat #26074-15). Microscopy images were captured using a ZEISS Axio Zoom.V15 microscope using the PlanNeoFluar Z 2.3x/0.57 FWD 10.6mm objective lens with a magnification of 260X. Digital images were captured at 2584 × 1936 pixel resolution at 12 bit/channel.

### Fluorescent in situ hybridization (sRNA-FISH)

sRNA-FISH experiment was carried out as described before^45^. Briefly, fresh unopened buds of strawberry were dissected and fixed in a 20 ml glass vial using 4% paraformaldehyde in 1xPHEM buffer (5 mM HEPES, 60 mM PIPES, 10 mM EGTA, 2 mM MgSO_4_ (pH 7). Fixation was done in a vacuum chamber at 0.08 MPa for 3 times, 15 min each. After fixation, samples were sent for paraffin embedding at histology lab from Nemours/Alfred I. duPont Hospital for Children (Wilmington, DE). Samples were sectioned using a paraffin microtome and dried on poly-l-lysine treated coverslips. Fluorescent in situ hybridization was modified from the protocol by Javelle et al. ^46^ by replacing the antibody with primary anti-Digoxigenin Fab fragment (Sigma-Aldrich cat# 11214667001) and secondary donkey anti-sheep IgG (H+L) AF647, AF568 or AF633 (Thermo Fisher Scientific cat# A-21448, A-21099, and A-21100). Briefly, samples were de-paraffin using Histo-Clear (Fisher scientific, 50-899-90147) and re-hydrate by going through an ethanol series of 95, 80, 70, 50, 30, 10% (vol/vol) (30 sec each) and water (1 min) at room temperature. After protease (Sigma, P5147) digestion (20 min, 37°C), samples were treated with 0.2% glycine (Sigma-Aldrich, cat # G8898) for 2 min. After two washes in 1xPBS buffer, samples were de-hydrated and then hybridized with probes overnight at 53.3°C. 10 ml of hybridization buffer contains 875 μl of nuclease-free H_2_O, 1.25 ml in situ hybridization salts, 5 ml of deionized formamide, 2.5 ml of 50% (wt/vol) dextran sulfate, 250 μl of 50x Denhardt’s solution, and 125 μl of 100 mg/ml tRNA. Hybridized slides were then washed twice using 0.2x SSC buffer (saline-sodium citrate). To immobilize the hybridized probes, samples were incubated for 10 minutes in freshly prepared EDC solution containing 0.13 M 1-methylimidazole, 300 nM NaCl (pH 8.0). Then samples were incubated for 1 hour and 15 minutes in 0.16 M N-(3-Dimethylaminopropyl)-N’-ethylcarbodiimide hydrochloride (EDC) (item 03450, Sigma-Aldrich, St. Louis, MO) solution. Afterward, samples were neutralized in 0.5% (w/v) glycine and blocked in 1x blocking buffer (1% blocking reagent in 1xTBS buffer), and 1x washing buffer (1% wt/vol BSA; Sigma-Aldrich-Aldrich, A7906) and 0.3% Triton x-100 in 1xTBS buffer) for 1 hour each. Samples were then incubated with primary antibody overnight at 4°C followed by 4x washes in 1x washing buffer, 15 min each. Samples were then incubated with a secondary antibody overnight at 4°C followed by 4x washes in 1x washing buffer, 15 min each. After final wash in 1xTBS buffer, samples were mounted using ProLong™ Glass Antifade Mountant (ThermoFisher Scientific, P36980). Antibodies were also hybridized to non-labeled samples as negative controls. Probes are shown in Supplementary Data 10.

### smFISH for phasiRNAs

Fifty probes were designed, corresponding to the top 50 21-nt phasiRNAs based on abundance in our libraries (i.e. the most abundant were selected). Each probe is 17 to 22 nt in length. The probes are designed using a web program: Stellaris Probe Designer (https://www.biosearchtech.com/support/tools/design-software/stellaris-probe-designer). The probes were ordered from LGC Biosearch Technologies with a 3’ end amino group and coupled with fluorophores TMR manually^47^. Fresh, unopened strawberry buds were prepared, embedded, sectioned the same way as the sRNA-FISH. Sample slides were then hybridized with smFISH probes at concentration 5 ng/ul in a 37°C hybridization oven overnight. After hybridization, the slides were washed 3x with 100 ml washing buffer, 20 min each wash. Samples were then washed with 2x SSC buffer, and were equilibrated for 2 min. Samples were mounted using ProLong™ Glass Antifade Mountant (ThermoFisher Scientific, P36980). smFISH probes for human AR mRNA were used as negative control. Probes are shown in Supplementary Data 10.

### Image acquisition

Spectral imaging was conducted on a Carl Zeiss LSM 880 laser scanning microscopy capable of both LSCM and multiphoton microscopy. The Zen software (v2.3; Carl Zeiss) was used for both acquisition of spectral images and linear spectral unmixing. Spectral data for pure Alexa Fluor^®^ fluorophores were used as positive controls, and non-labeled samples were used to obtain autofluorescence spectra for linear spectral unmixing. Brightness and contrast of images in the same figure panel were adjusted equally and linearly in Zen software (Carl Zeiss).

## Data availability

All the sRNA and RNAseq data generated during this study are under submission to NCBI’s SRA (Sequence Read Archive). All the RNAseq and sRNA data from public databases used for the analysis are listed in Supplementary Data 11.

## Author Contributions

B.C.M. and S.P. designed the research; S.P. performed the experiments and analyzed the data; K.H. performed fluorescent *in situ* hybridizations, imaging analysis; S.B. helped with staging of buds/anthers; J.L.C. supervised *in situ* hybridizations; E.M.K. helped with collecting buds/anthers of columbine. S.P. and B.C.M wrote and revised the manuscript with input from all co-authors.

## Acknowledgements

We thank members of the Meyers lab for helpful discussions, and Joanna Friesner for assistance with editing. We thank Mayumi Nakano for assistance with data handling. This work was supported by resources from the Donald Danforth Plant Science Center and the University of Missouri – Columbia; additional support was provided by NSF Plant Genome Research Program Award 1754097 and USDA/NIFA Award 2019-67013-29010. Microscopy equipment was acquired with a shared instrumentation grant (S10 OD016361) and access was supported by the NIH-NIGMS (P20 GM103446), the NSF (IIA-1301765) and the State of Delaware.

