## Supplemental Figures for "Pre-meiotic, 21-nucleotide Reproductive PhasiRNAs Emerged in Seed Plants and Diversified in Flowering Plants"

**Supplemental Figure 1. miR11308 and 21-nt phasiRNA pathway in wild strawberry**

A. A polycistronic precursor cluster of miR11308b and miR11308c encoded on chromosome 6 (coordinates indicated above the structure). The mature miR11308 and miR11308* sequences are marked in blue and yellow respectively.

B. Two miR11308b and miR11308c precursors are transcribed into a single transcript. sRNA and RNA-seq data are displayed in the Integrative Genomic Viewer (IGV).

C. Genome-wide distribution of miR11308 and miR2118/482 family members (inner circle, red dots) and their 21-PHAS loci (outer cycle, green dots). The outermost cycle represents chromosomes from 1 to 7, plus chromosome "8" which represents an amalgam of the unassembled regions. Red colored loci are reproductive-enriched 21-*PHAS* loci triggered by miR11308 family members.

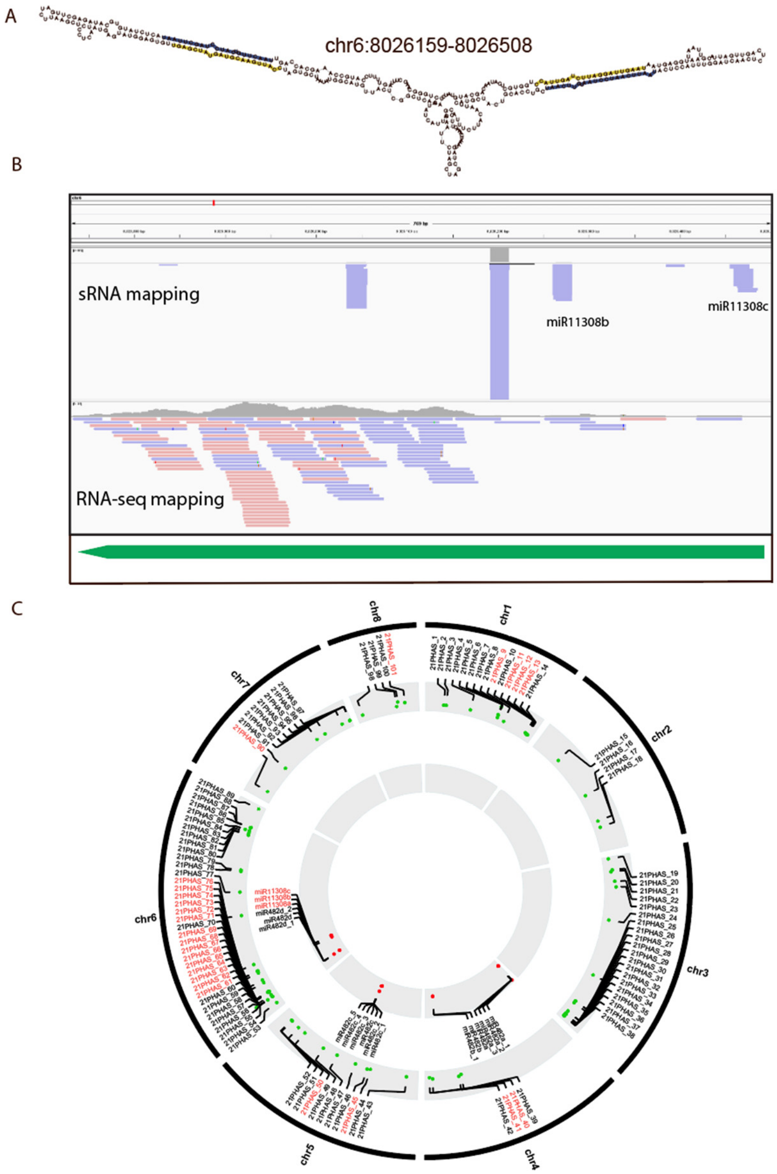

**Supplemental Figure 2. miR2118/482 family and 21-nt phasiRNA pathway in wild strawberry**

A. The abundance of miR2118/482-triggered 21-nt phasiRNAs across different tissues in wild strawberry. The key at right shows the abundance in unit of log2(RP20M). The lowermost 11 loci are reproductive-enriched (fold change >1.5) while the lowermost six loci are non-coding.

B. Abundance of miR2118/482 variants in different tissues in wild strawberry.

C. Alignment of members of the miR2118/482 family in wild strawberry. The degree of conservation is represented by intensity of blue color, and the consensus sequence of the alignment is shown with a sequence logo.

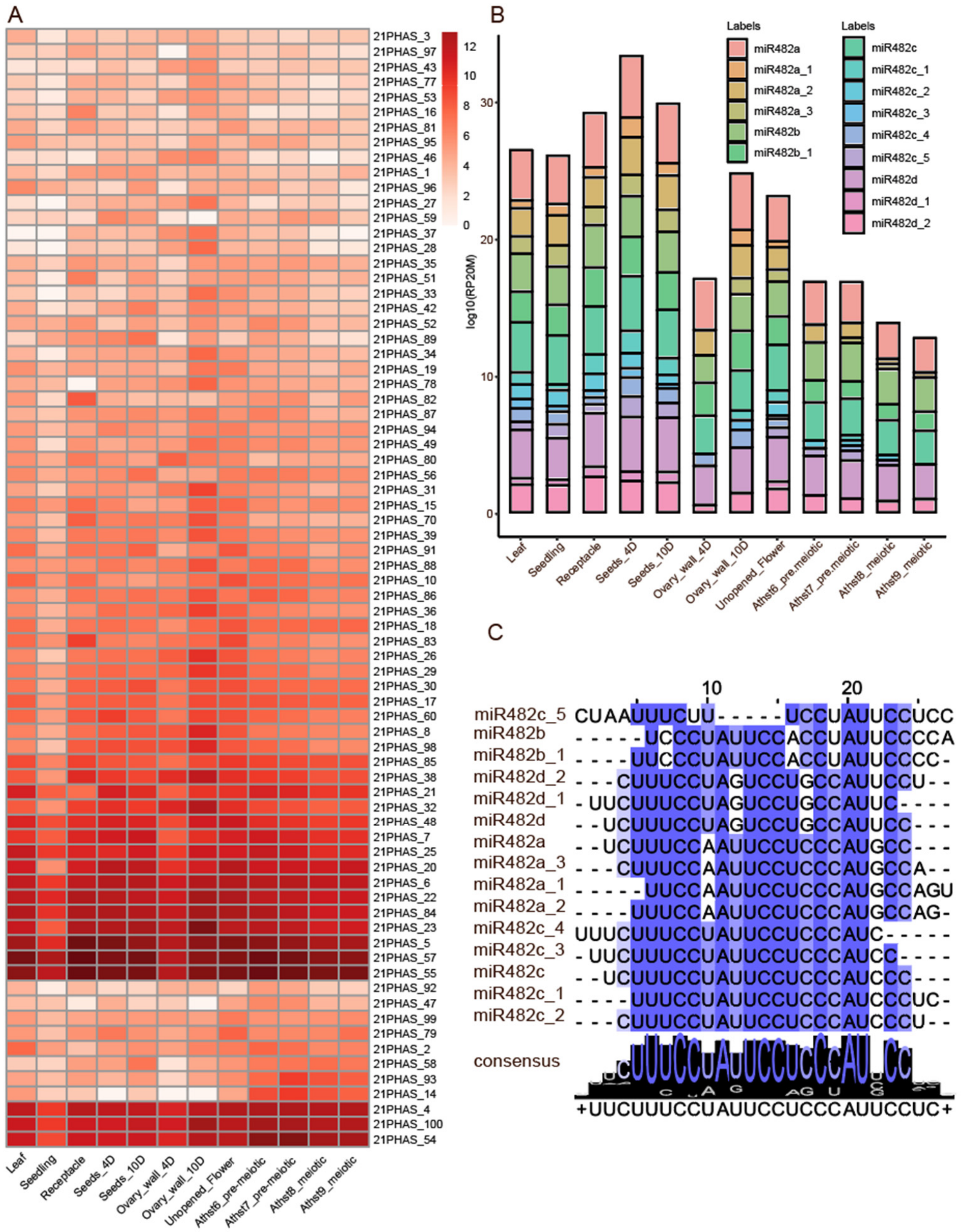

**Supplemental Figure 3. Schematic diagram of smFISH of phasiRNAs plus the miR2118/482 family and 21-nt phasiRNA pathway in rose.**

A. Schematic diagram of smFISH using a pool of 50 abundant phasiRNAs in wild strawberry, yielding the data shown in Figure 3. E: epidermis, En:endodermis, M: middle layer, T: tapetum, MMC: microspore mother cell.

B. Bud stages of rose from which anthers were harvested. Bst indicates bud stage, anth indicates anther, stage 1: 1 cm buds, stage 2: 1.25 cm buds, stage 3: 1.5 cm buds, Stage 4: 2 cm buds.

C. The abundance of miR2118/482-triggered 21-nt phasiRNAs abundance among different tissues in rose.

D. Alignment of members of the miR2118/482 family in rose. The degree of conservation is represented by the intensity of the blue color and consensus sequence of the alignment is shown with a sequence logo.

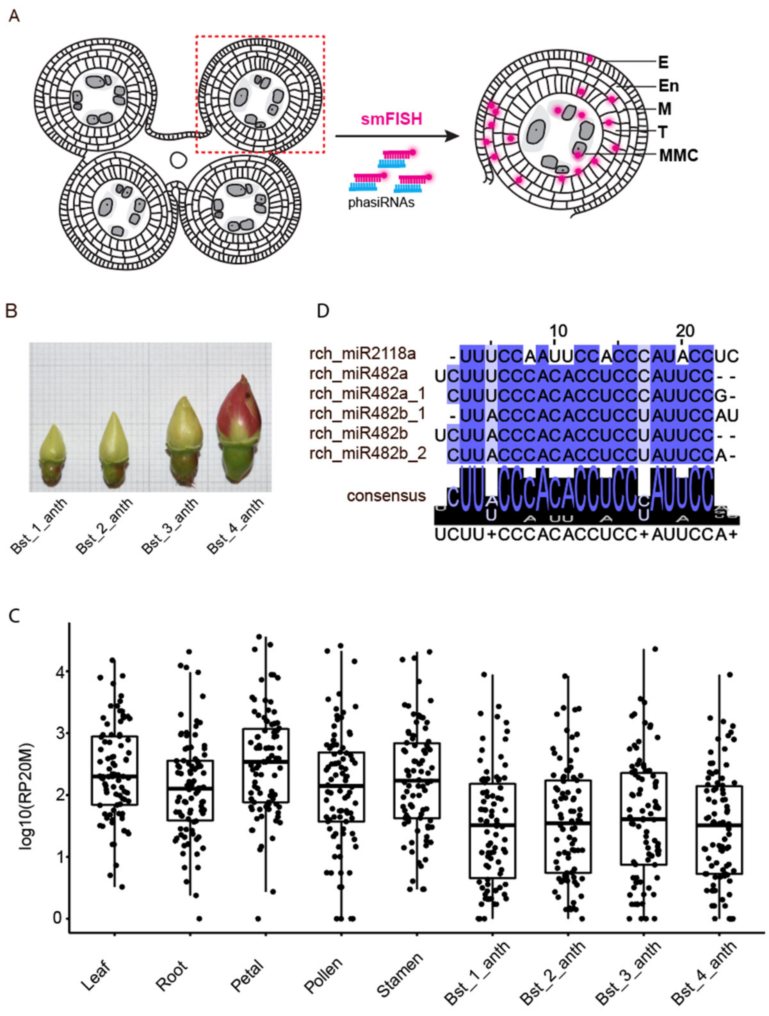

**Supplemental Figure 4. The 21-nt phasiRNA pathway in columbine**

A. Bud stages in columbine, Bst indicates bud stage, anth indicates anther, stage 1: 1 to 2 mm buds, stage 2: 3 to 5 mm buds, stage 3: 6 to 10 mm buds, stage 4: 10 to 20 mm buds.

B. The abundance of miR2118/482- and aco-cand81-triggered 21-nt phasiRNAs in vegetative/reproductive tissues in columbine. The key at right displays the abundance in units of log2(RP20M).

C. Above: Sequence logo denoting conservation of the target site of aco_cand81, determined from 65 21-*PHAS* loci. Below: Nucleotide sequence alignment of 21-*PHAS* loci with sequence similarity denoted by the intensity of the blue color, showing that the aco_cand81 target site is the only conserved region for all the precursors.

D. A polycistronic precursor cluster of aco_cand81e and aco_cand81f in chromosome 4. The mature aco_cand81 and aco_cand81* sequences are marked in blue and yellow respectively.

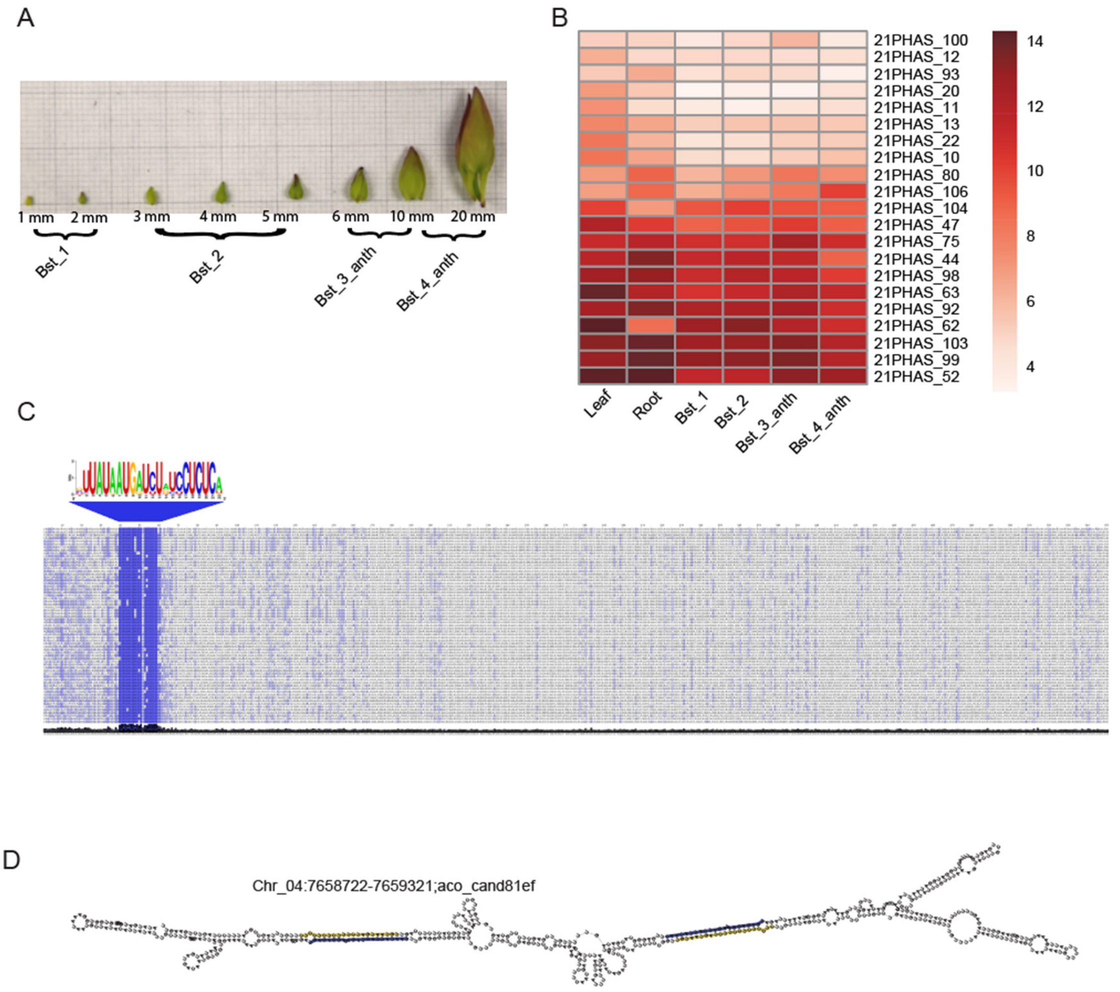

**Supplemental Figure 5. Target site conservation of miR2118/482 derived 21-PHAS loci in columbine and polycistronic precursors of miR2118/482**

A. Above: Sequence logo denoting conservation of the target site of aco_miR2118/482 for 47 21-*PHAS* loci. Below: Nucleotide sequence alignment of 21-*PHAS* loci with sequence similarity denoted by the intensity of blue color showing that aco_miR2118/482 target site is the only conserved region for all the precursors.

B. Two polycistronic precursor clusters of aco_miR2118/482 four variants in chromosome 3. The mature aco_miR2118/482 and aco_miR2118/482* sequences are marked in blue and yellow respectively.

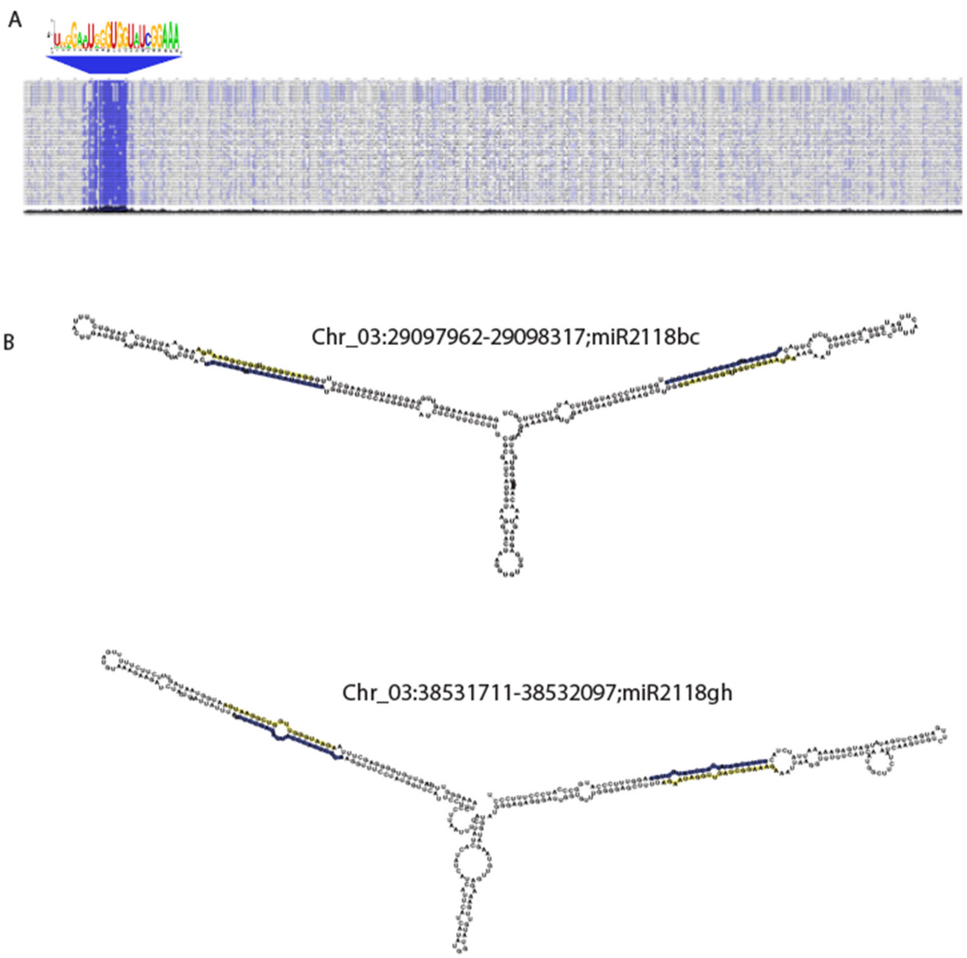

**Supplemental Figure 6. Genome-wide distribution of triggers and 21-PHAS Loci in columbine**

Genome-wide distribution of aco_cand81 and miR2118/482 family members and their 21-*PHAS* loci represented by inner (red) and outer (green) cycles respectively.

A

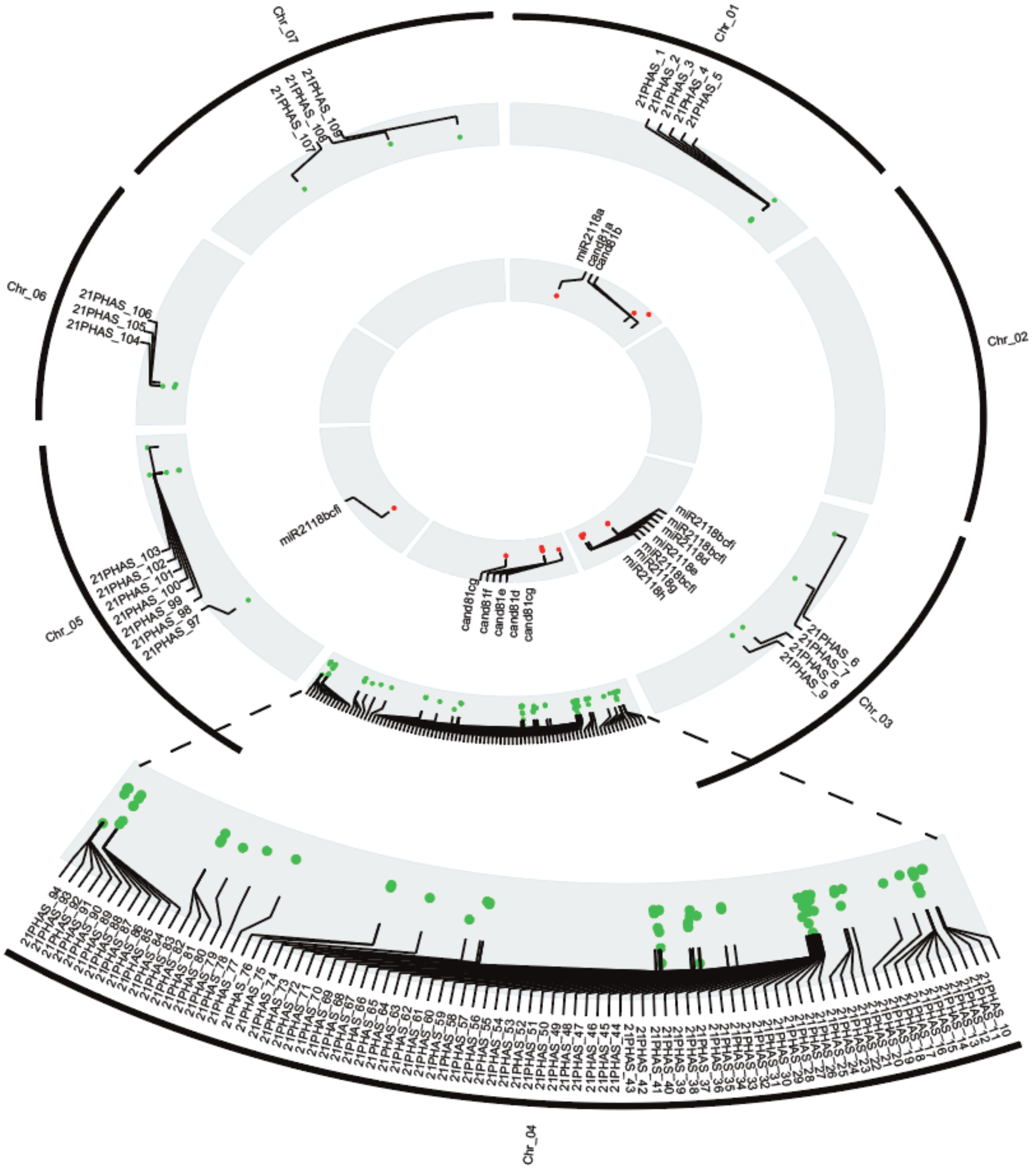

**Supplemental Figure 7. Transverse sections of anthers from 2-20 mm buds in size covering four different stages in columbine.**

Anthers were fixed in a FAA solution and embedded using the Quetol epoxy resin, sectioning at 0.5 µm and stained using 0.05% toluidine blue O. Black scale bars correspond to 50 *µm*.

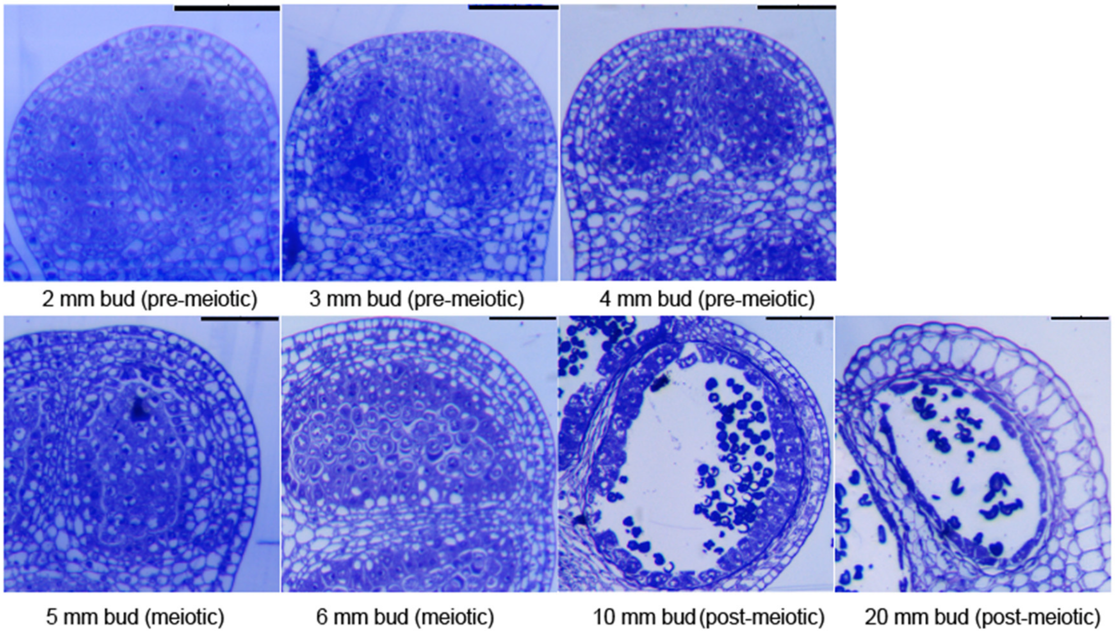

**Supplemental Figure 8. Reproductive 21-nt phasiRNAs in flax and their trigger miR2118/482**

A. Bud stages in flax, Bst indicates the bud stage, stage 1: 3 to 5 mm buds, stage 2: 5 to 8 mm buds, stage 3: 8 to 10 mm buds.

B. Accumulation of 21-nt phasiRNAs in different tissues in flax. The key at right indicates the abundance in unit of log2(RP20M).

C. Alignment of variants of miR2118/482 family in flax.

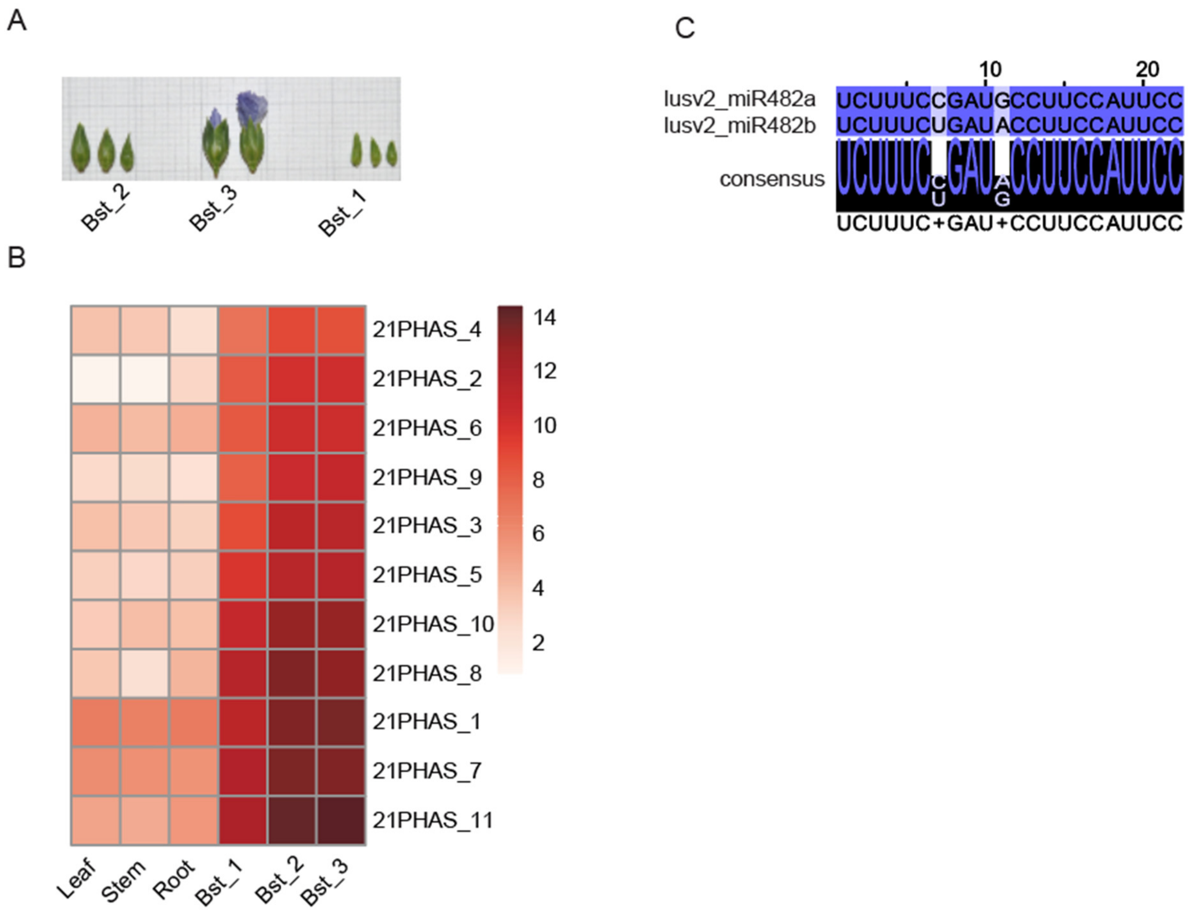

**Supplemental Figure 9. Protein diversity of RNA-dependent RNA polymerase (RDR) and Dicer-like (DCL) family members encoded in genomes of wild strawberry, rose, columbine, flax, other representative species from gymnosperms, eudicots, and monocots.**

A. Phylogenetic tree of RDR family members of wild strawberry (Fv), rose (Rc), columbine (Ac), flax (Lu), norway spruce (Pa), ginko (Gb), *Amborella* (Atr), soybean (Gm), tomato (Sl), *Arabidopsis* (At), *Asparagus* (Ao), maize (Zm), and rice (Os). The asterisk indicates the species with duplicated copies mentioned in the main text.

B. Phylogenetic tree of DCL family members of same species as A. The asterisk indicates the species with duplicated copies mentioned in the main text.

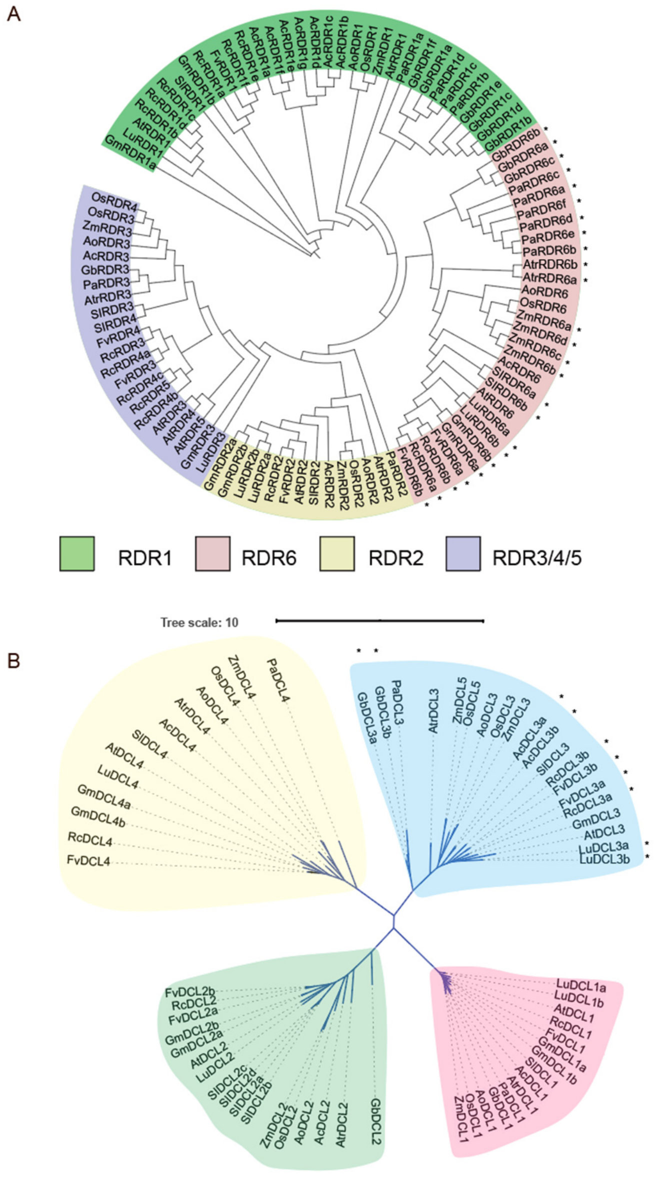
